# A comparison of Remdesivir versus Au_22_Glutathione_18_ in COVID-19 golden hamsters: a better therapeutic outcome of gold compound

**DOI:** 10.1101/2020.10.16.342097

**Authors:** Zhesheng He, Fei Ye, Chunyu Zhang, Jiadong Fan, Zhongying Du, Wencong Zhao, Qing Yuan, Wenchao Niu, Fuping Gao, Bo He, Peng Cao, Lina Zhao, Xuejiao Gao, Xingfa Gao, Bo Sun, Yuhui Dong, Jincun Zhao, Jianxun Qi, Huaidong Jiang, Yong Gong, Wenjie Tan, Xueyun Gao

## Abstract

We firstly disclose single compound yields better therapeutic outcome than Remdesivir in COVID-19 hamster treatments as it is armed with direct inhibition viral replication and intrinsic suppression inflammatory cytokines expression. Crystal data reveals that Au (I), released from Au_22_Glutathione_18_ (GA), covalently binds thiolate of Cys145 of SARS-CoV-2 M^pro^. GA directly decreases SARS-CoV-2 viral replication (EC50: ~0.24 μM) and intrinsically down-regulates NFκB pathway therefore significantly inhibiting expression of inflammatory cytokines in cells. The lung viral load and inflammatory cytokines in GA-treated COVID-19 transgenic mice are found to be significantly lower than that of control mice. When COVID-19 golden hamsters are treated by GA, the lung inflammatory cytokines levels are significantly lower than that of Remdesivir while their lung viral load are decreased to same level. The pathological results show that GA treatment significantly reduce lung inflammatory injuries when compared to that of Remdesivir-treated COVID-19 golden hamsters.

**One Sentence Summary:** We found that gold cluster molecule directly inhibits SARS-CoV-2 replication and intrinsically suppresses inflammatory cytokines expression in COVID-19 transgenic mouse and golden hamster model, gold cluster providing a better lung injury protection than Remdesivir in COVID-19 golden hamsters via intranasally dropping administration.

## Introduction

To date, over 4 Million people have succumbed to COVID-19 infections worldwide, with little sign of this global pandemic being swiftly brought under control. Despite tremendous global efforts in identifying a suitable drug against COVID-19, no drug has been proven to effectively treat COVID-19 infections. In comparison of biological agents, chemical drugs are of unique advantages in dealing with the COVID-19 pandemic: they are easily produced in large scale with low cost, thus satisfying the huge number of COVID-19 patients in low income countries. Importantly, chemicals allow for efficient handling, storing and distributing to patients living in environments unsuitable for biological agents. Several traditional chemicals are currently repurposed for COVID-19 treatment (1, 2, 3, 4, 5, 6, 7). For example, Remdesivir was shown to directly inhibit SARS-CoV-2 replication but failed to intrinsically suppress inflammatory cytokines expression in patients (3, 7). In comparison, Ruxolitinib or Acalabrutinib intrinsically suppressed inflammatory cytokines expression, but failed to directly inhibit virus replication (4, 6).

We firstly propose that single compound combination of directly inhibition viral replication and intrinsically suppression inflammatory cytokines expression should yield better therapeutic results in COVID-19 treatment. The SARS-CoV-2 M^pro^ has been used as critical drug target for COVID-19 treatment as it plays a key role in SARS-CoV-2 replication, and organic compounds have developed to yield a Michael adduct with Cys145 of the catalytic dyad of M^pro^ (8, 9). We speculated that Au (I) ion would yield a Michael adduct with Cys145 of M^pro^ as previous reports showed that the Au (I) ion, released from gold compound, inactivates *Echinococcus Granulosus* Thioredoxin Glutathione Reductase via covalently bind with Cys519 and Cys573 (10, 11). While gold compound, like Auranofin (AF), have previously been approved for Rheumatoid arthritis (RA) treatment as it well suppresses inflammatory cytokines of patients (12, 13), no such treatment is currently available for COVID-19 treatment *in vivo*. Here, we propose a novel compound, Au_22_Glutathione_18_, for COVID-19 treatment via directly inhibition of SARS-CoV-2 replication and intrinsically suppression of inflammation cytokines expression. We set out to investigate if this gold cluster possesses therapeutic properties in mitigating the effects of COVID-19 infections.

## Results

### Gold compound (AF or GA) firstly associates with the catalytic domain of SARS-CoV-2 M^pro^ and finally produce Au-M^pro^ adduct

The gold cluster was synthesized by a chemical method (**SI 1**). According mass spectra studies and density functional theory (DFT) simulations, the Au_22_Glutathione_18_ (GA) molecule formula is proposed in **Figure S1**. We firstly study if gold compound (AF or GA) associates with hydrophobic domain surround the catalytic dyad of M^pro^ via molecular dynamic (MD) simulations (**SI 2**). In this theory study, the crystal structure of M^pro^ was obtained from the PDB database (PDB 6LU7). The AF and GA molecular structure were performed by using GaussView 5.0 and Gaussian 09 software packages. As shown in **Figure 1A**, there is one hydrophobic cavity with its volume over 500 Å^3^ in each M^pro^ monomer, which surrounds the catalytic site of Cys145 (green or purple region of M^pro^ monomer). In this work, we utilized AF or GA as ligand to complex with M^pro^. For the AF-M^pro^ or GA-M^pro^ complexing system, we studied the root-mean-square deviation (RMSD) to learn the binding stability. Obviously, the AF-M^pro^ or GA-M^pro^ reached binding equilibrium at about 10 ns, and the binding conformation remained stable afterwards (**Figure S2**). The average binding energy of AF-M^pro^ and GA-M^pro^ was ~50.47 kcal/mol. and ~722.94 kcal/mol., respectively (**Figure S3**). The GA (yellow) and AF (orange) bind to the hydrophobic cavity around the Cys145, and this makes Au atom very close to S of Cys145 (**Figure 1B**). In AF-M^pro^ system, Van del Vaals force plays main role to keep system stable where the Glu166 and Gln189 of M^pro^ interact with AF via hydrogen bonds. While the electrostatic force contributes the main interaction in GA-M^pro^ system in **Figure 1B**, Arg4, Arg40, Lys137 and Arg188 of M^pro^ formed four salt bridges with glutathione of GA (with yellow S atoms).

**Fig. 1.**
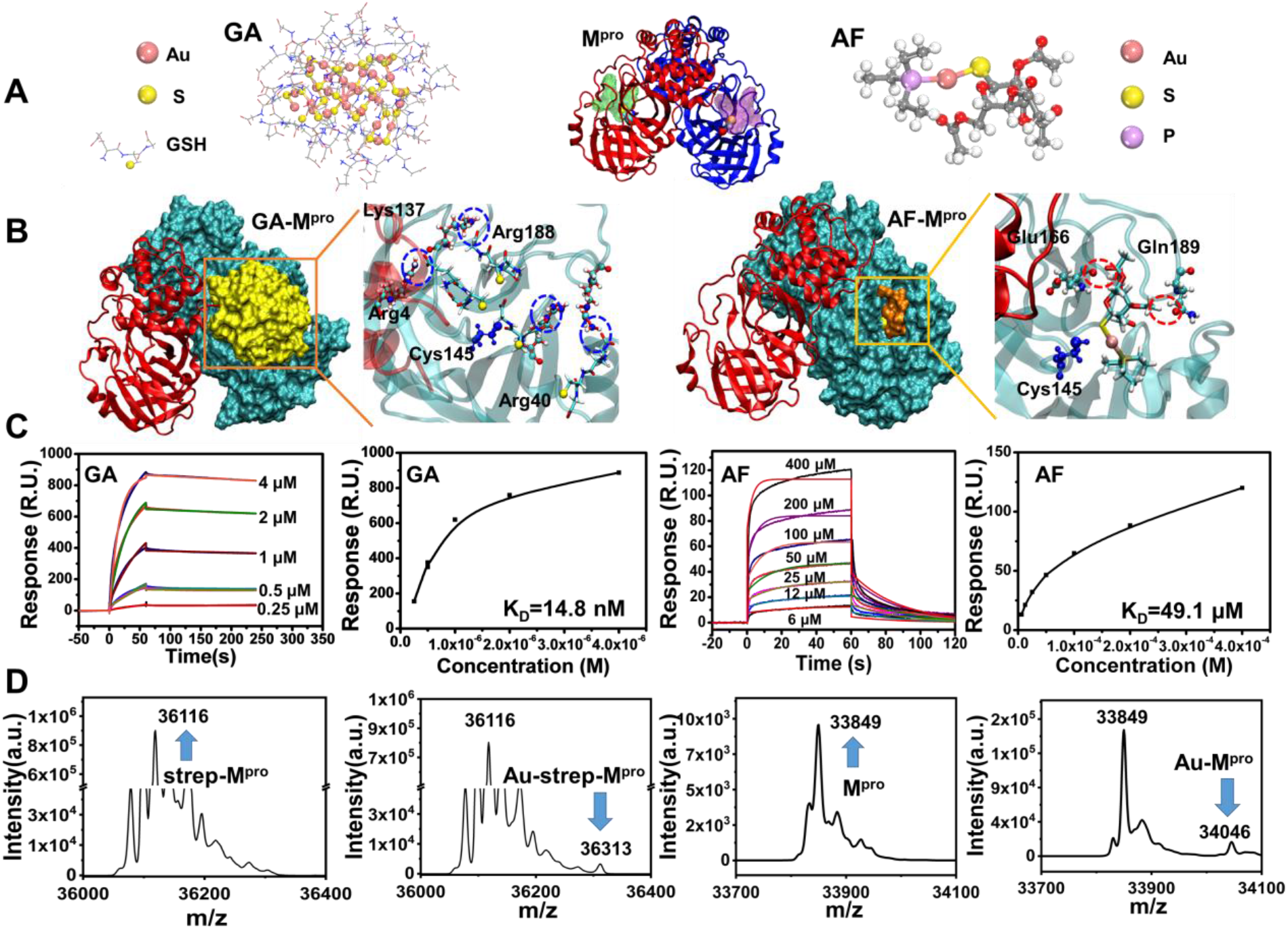
Theory simulations and SPR studies of gold compound complex with M^pro^ and ESI-Mass spectra of final Au-M^pro^ adduct. (**A**) The molecular structure of GA, AF, and SARS-CoV-2 M^pro^. (**B**) Molecular dynamic simulations of GA and AF complex with M^pro^. Both compounds stably bind the hydrophobic domain surround the catalytic dyad of M^pro^, GA-M^pro^ (binding energy ~722.94 kcal/mol.) is more stable than AF-M^pro^ (binding energy ~50.47kcal/mol.). (**C**) SPR studies of GA and AF association with M^pro^, K_D_ of GA is ~ 14.8 nM and that of AF is ~ 49.1 μM. (**D**) ESI-Mass spectra of Au-M^pro^ adduct which is extracted from GA treated HEK293F cells (**left**), ESI-Mass spectra of Au-M^pro^ adduct which is purified from AF incubated M^pro^ solution (**right**). Note that molecule weight of strep-M^pro^ is 36116 Da and Apo M^pro^ molecular weight is 33849 Da. Mass data revealed one Au ion bind one M^pro^ monomer.

The experimental association affinity between gold compound (AF or GA) and M^pro^ was further studied by surface plasma resonance (SPR) method **(SI 3)**. After M^pro^ was fixed in chip, serial dose of gold compound (AF or GA) was introduced into solution and the time dependent optical signal were tracked. As shown in **Figure 1C**, the K_D_ of GA-M^pro^ and AF-M^pro^ is ~14.8 nM and ~49.1 μM, respectively. This SPR data implied that GA is with stronger affinity with M^pro^ when compared with that of AF, and this experimental result matches the aforementioned molecular dynamic simulations where the average binding energy of AF-M^pro^ was ~50.47 kcal/mol. and that of GA-M^pro^ is the ~722.94 kcal/mol.. Although theory and SPR studies revealed gold compound can tightly bind the M^pro^, we do not know the final product form of GA-M^pro^ and AF-M^pro^. To clarify this issue, strep tagged-M^pro^ (strep-M^pro^) were expressed in HEK293F cell and high concentration of GA were introduced in cell culture media, this would produce final Au-M^pro^ adduct in cytoplasm (**SI 4**). AF were directly introduced to none strep tagged-M^pro^ (M^pro^ in Apo form) solution to get final Au-M^pro^ adducts as HEK293F cell could not bear toxicity of high concentration AF in cell culture media (**SI 4**). After gold compound treated M^pro^ was purified, ESI mass spectra was used to check the final product of GA-M^pro^ or AF-M^pro^. In **Figure 1D**, both GA and AF treated M^pro^ finally produce Au-M^pro^ adduct where one Au atom is added to one M^pro^. These studies verified that GA or AF firstly associated with M^pro^ and finally produce Au-M^pro^ adduct in one Au atom per one M^pro^ manner.

### Au (I), released from gold compound, covalently binds the thiolate of Cys145 of M^pro^ via experimental crystal structure studies

In order to examine molecular structural basis of gold compound inactive M^pro^ in *vitro/vivo*, we determined the SARS-CoV-2 M^pro^ experimental crystal structures in the M^pro^ crystal incubated with AF (AF incubating form), M^pro^ crystal incubated with GA (GA incubating form), and M^pro^ crystal only (the native form), see details in **SI 5, Figure S4, and Table S1**. **In Figure 2A**, the M^pro^ molecular structures treated with AF or GA are highly similar, and share most features of the crystal structures of the apo SARS-CoV-2 M^pro^ determined recently (8, 9). However, crystal structural analysis showed that the densities of two Au (I) ions were found clearly to be very close to the thiol residues of Cys145 and Cys156 (**Figure 2B**). The position of two Au (I) ions was confirmed by applying the anomalous difference Fourier maps, two Au (I) ions are defined as Au(I) 1 and Au (I) 2, respectively. In **Figure 2C and Figure S5**, the Au-S bond length is 2.3Å in M^pro^, such short bond length confirms that AF or GA can release Au (I) ions which covalently bind to the thiolate of Cys145 and Cys156 of M^pro^. The thiolate of Cys519 and Cys573 residues of *Echinococcus Granulosus* enzyme covalently interacts with Au (I) ions were previously reported (10), which supports Au (I) irreversibly interact with thiolate of Cys145 of the catalytic dyad of M^pro^ in this study. Further temperature factor analysis shows that the occupancy of Au (I) ion is partial and the occupancy factors is about 33% for Cys145 and 11% for Cys156, indicating that the Au (I) ions released from AF or GA are gradually bind the thiolate of Cys145 and Cys156 of M^pro^ molecules in solution.

**Fig. 2.**
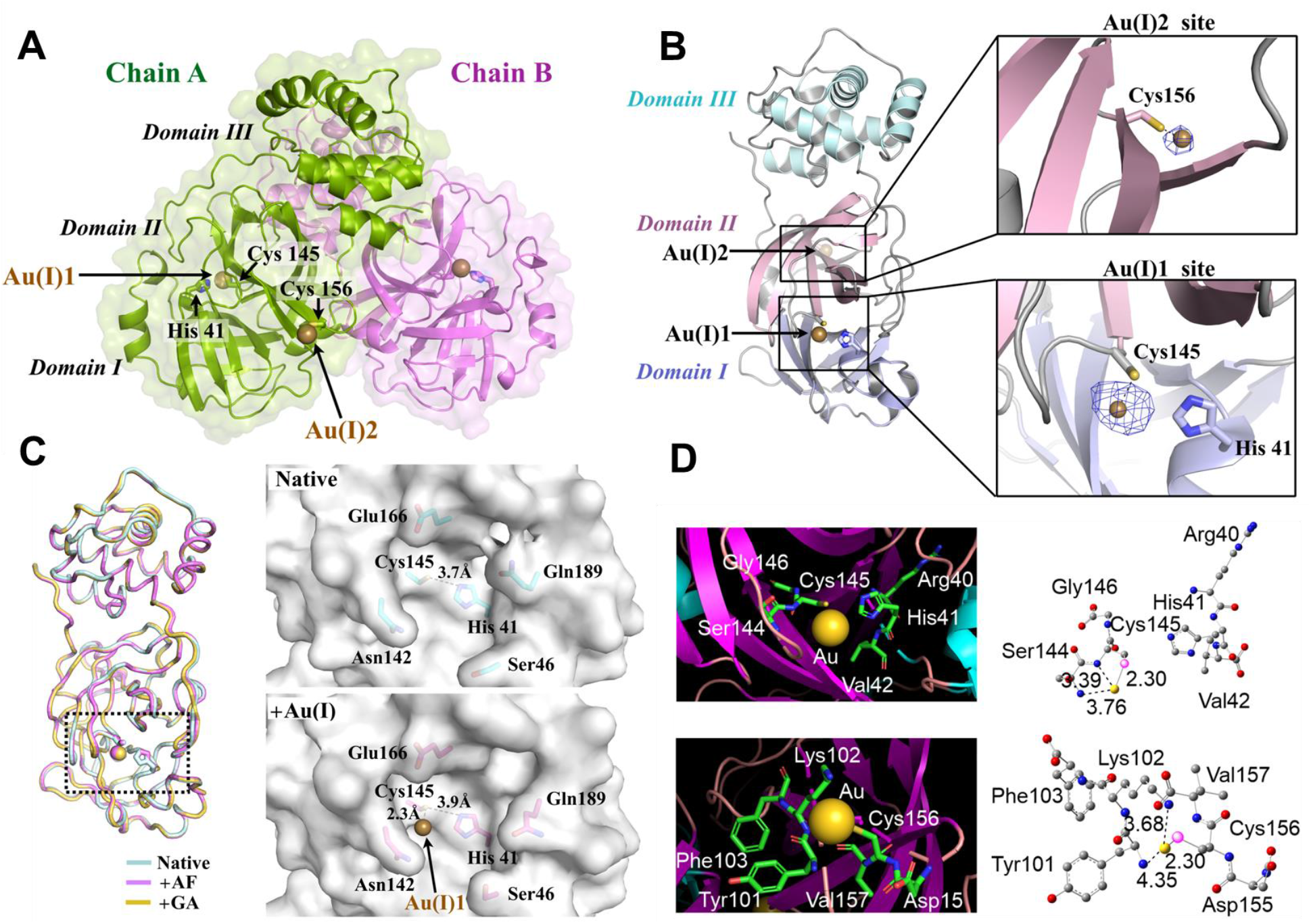
The X-ray crystal structure of AF and GA treated M^pro^ in Au-S covalent bind state. (**A**) Surface presentation of the M^pro^ homodimer in Au-S bound state with Chain A and Chain B shown in green and violet, respectively. (**B**) The presentation of one M^pro^ monomer with Domain I-III shown in light blue, light pink and pale cyan. Enlarged views of the Au (I)-S bound sites. Anomalous difference Fourier maps (blue mesh, contoured at 5 sigma) are shown for Au (I) 1 and Au (I) 2. Residues His41, Cys145 and Cys156 are shown in sticks and two Au (I) ions are shown in spheres. (**C**) Comparison of Au (I)-Cys145 bound state with the native state of M^pro^. Superposition of crystal structures of AF treated (purple), GA treated (yellow) and native M^pro^ (blue). The catalytic pocket of native and Au (I)-S bound M^pro^ in surface presentation and the surrounding residues shown in sticks. (**D**) DFT calculation of interaction between Au (I) ions and Cys145 and Cys156 of M^pro^, respectively. Left panels show protein catalytic pockets consisting of amino acids and one Au (I) ion. Right panels represent geometrically relaxed structures for the catalytic pockets encapsulating the Au (I), all Au−N atomic distances (in Å) within 5 Å are labeled with the corresponding distances. C, N, O, S, and Au atoms are displayed in grey, blue, red, pink, and yellow, respectively.

Although the M^pro^ monomer contains 12 Cys residues (Cys16, Cys22, Cys38, Cys44, Cys85, Cys117, Cys128, Cys145, Cys156, Cys160, Cys265, Cys300), only Cys145 and Cys156 specifically bind to the Au (I). To further verify the covalently binding of Au (I) to the S atom of Cys145 and Cys156, we calculated the interaction energies between Au and M^pro^ protein using density functional theory (**SI 6**) method. Our analysis showed that the bond dissociation energies (*E*_BD_’s) between Au (I) and Cys145 are ~ 46.1 kcal/mol. and that of Au (I) and Cys156 are ~ 26.5 kcal/mol. In (**Figure 2D**. The larger *E*_BD_ value strongly suggests that the Au ion covalently bind to Cys145 and lock the active pocket of M^pro^, thus efficiently inhibiting catalytic activity.

### Gold compound (AF or GA) inhibits SARS-CoV-2 M^pro^ activity, suppress SARS-CoV-2 replication, and block inflammatory cytokines expression in cell assays

To exam whether AF or GA effectively inhibits M^pro^ activity, we determined the IC50 of AF or GA using a previously reported method (14). M^pro^ activity was measured using a fluorescence resonance energy transfer (FRET) assay. To this end, a fluorescence labeled substrate, (EDNAS-Glu)-Ser-Ala-Thr-Leu-Gln-Ser-Gly-Leu-Ala-(Lys-DABCYL)-Ser, derived from the auto-cleavage sequence of the viral protease was chemically modified for enzyme activity assay (**SI 7**). As shown in **Figure 3A**, the IC50 of AF was ~0.46 μM, and the IC50 of GA was ~0.11 μM.

**Fig. 3.**
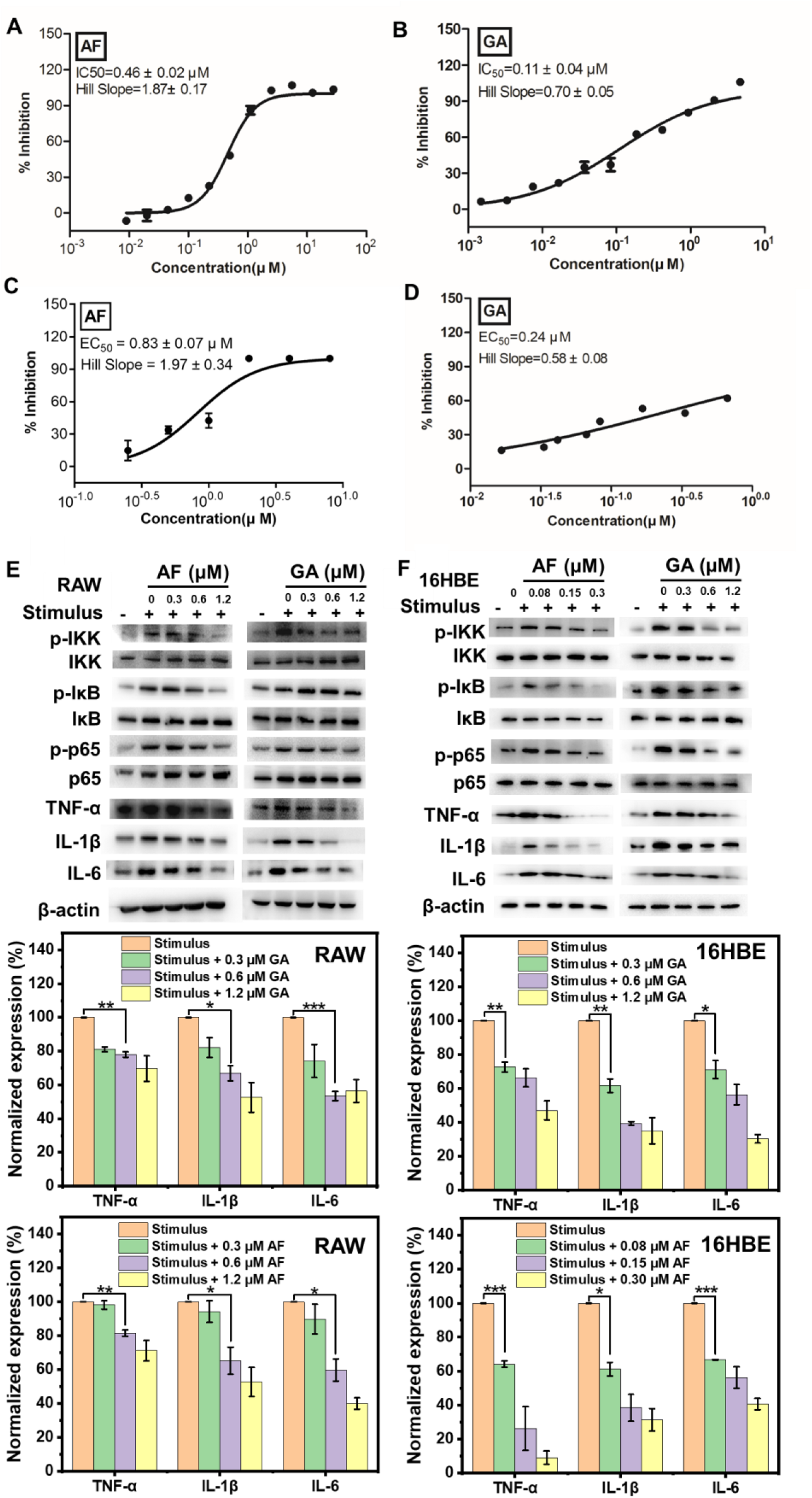
AF and GA inhibit M^pro^ activity, suppress SARS-CoV-2 replication, and inactivate the NFκB pathway and suppress inflammatory cytokines expression in cells. (**A**) IC50 of AF is ~ 0.46 μM. (**B**) IC50 of GA is ~ 0.11 μM. (**C**) EC50 of AF is ~ 0.83 μM. (**D**) EC50 of GA is ~ 0.24 μM. (**E**) Low dose of AF (0.6 μM) and GA (0.6μM) significantly inhibit IL-6, IL-1β, TNF-α inflammatory cytokines expression in RAW macrophages (unpaired t-test, ***p<0.001, **p <0.01, *p <0.05). (**F**) Low dose of AF (0.08μM) and GA (0.3 μM) significantly suppress NFκB activation, thus inhibiting IL-6, IL-1β, TNF-α inflammatory cytokine expression in human 16HBE bronchial epithelial cells (unpaired t-test, ***p<0.001, **p <0.01, *p <0.05).

To assess whether GA inhibits M^pro^ activity in mammalian cells, we transiently transfected HEK293F cells with a plasmid of strep-tagged SARS-CoV-2 M^pro^ gene. The M^pro^ gene was expressed 24 hours in HEK293F cells, then GA was added to the culture medium at a final concentration of 500 μM, and cell cultured for an additional 24 hrs. After cell harvest, SARS-CoV-2 M^pro^ extracted from GA-treated HEK293F cells was purified and analyzed for enzyme activity (**SI 7**). As shown in **Figure S6**, the GA treated-SARS-CoV-2 M^pro^ activity was reduced to approx. 60% of control SARS-CoV-2 M^pro^. ICP-MASS results indicated that when M^pro^ was expressed in GA-treated mammalian cells, gold can bound to SARS-CoV-2 M^pro^ (**Figure S6**). We failed to obtain similar results for AF in cell assays as AF exhibited strong cytotoxicity when HEK293F cell cultured with AF.

We measured the EC50 of AF or GA to evaluate if they inhibit SARS-CoV-2 replication in Vero cells, using a recently described method (5) (**SI 7**). As shown in **Figure 3B**, the EC50 of AF was approx. 0.83 μM and EC50 of GA approx. 0.24 μM. However, for Vero cell the CC50 value of AF is ~ 2.27 μM and that of GA approx. 1.11mM (**Figure S7**). The therapeutic index (CC50/EC50) of AF is 2.73 and that of GA is 4,625, this implied GA has better potential in COVID-19 treatment.

COVID-19 viral infections are characterized by infections of bronchial epithelial cells, resulting in activation of inflammatory cytokine gene expression via the NFκB pathway, and these cytokines will in turn activate macrophages, which as a result acquire an inflammatory status to profoundly produce inflammatory cytokines (15, 16). Recent reports revealed that among versatile inflammatory cytokines, the IL-6, IL-1β, TNF-α play key roles in inflammation development of modest and severe infections (16, 17). We firstly assessed whether GA or AF inactivate the NFκB pathway and suppress inflammatory cytokine expression levels in inflammatory human 16HBE bronchial epithelial cells (**SI 7**). To test whether AF or GA inactivate the NFκB pathway in macrophage cells, and as a result down-regulate expression of IL-6, IL-1β, TNF-α, RAW264.7 macrophage was incubated with AF or GA at different concentrations for 24 hours. As shown in **Figure 3C**, the low dose of AF (0.6 μM) or GA (0.6 μM) significantly suppressed IL-6, IL-1β and TNF-α expression levels in RAW264.7 macrophage cells. For human bronchial epithelial cells, AF (0.08 μM) and GA (0.3μM) significantly inhibited phosphorylation of IKK, IκB, p65, thus significantly inhibiting IL-6, IL-1β, and TNF-α inflammatory cytokine expression. *Jue* et al. had demonstrated that the Cys179 of IKKβ plays a critical role in activation of NFκB pathway, and the anti-inflammatory activity of gold compound may depend on modification of thiolate of Cys179 via Au ion (18). For 16HBE cell the CC50 of AF is ~ 0.63 μM and that of GA approx. 1.06 mM, and for RAW cell the CC50 of AF is ~ 2.63 μM and that of GA approx. 1.44 mM (**Figure S7**). This implied that GA has better safety in COVID-19 treatment.

### GA inhibits SARS-CoV-2 replication and decreases inflammatory cytokine level in lungs, and protects lungs from inflammatory injury in COVID-19 transgenic mouse model

To evaluate the safety of administering AF or GA into COVID-19 animal model, we first tested the toxicity of AF and GA. The reported mice intraperitoneal LD50 for AF was approx. 33.8 mg/kg.bw (19), and that for GA more than 2000 mg/kg.bw (all mice survived, see **SI 8**). The SD rat intraperitoneal LD50 for AF was found to be approx. 25.5 mg/kg.bw (19), while that for GA approx. 576 mg/kg.bw (**SI 8)**. Together, these animal toxicity and aforementioned cell toxicity data (**Figure S8**) strongly suggested that GA is safer for mice/rats treatment than AF. We therefore continued investigation of GA in a COVID-19 transgenic mouse. To evaluate whether GA inhibit SARS-CoV-2 replication and suppress lung inflammation injury, the Ad5-hACE2-transduced mice were generated according to recently published methods (20). Briefly, eighteen BALB/c mice were randomly divided into three groups, namely GA (GA treated COVID-19 mice), NS (0.9% NaCl treated COVID-19 mice), Mock (0.9% NaCl treated mock infected-mice), all mice were anesthetized with pentasorbital sodium and transduced intranasally with 2.5×10^8^ FFU of Ad5-ACE2 in 50 μL DMEM (**SI 9**). Five days post transduction, the mice in GA group received a dose of 30 mg/kg.bw GA via intraperitoneal injection (i.p.). For mice in NS group, an equivalent volume of normal saline (0.9% NaCl) via intraperitoneal injection (i.p.) as vehicle. For mice in Mock group, an equivalent volume of normal saline (0.9% NaCl) via intraperitoneal injection (i.p.) without SARS-CoV-2 infection as a control. One hour after GA or normal saline treatment, mice in GA or NS group were infected intranasally using SARS-CoV-2 (1×10^5^ PFU) in a total volume of 50 μL DMEM. After virus infection at day 0, mice received GA or NS treatment for further three times as shown in **Figure 4A**. All mice were euthanized at day 4 post infection, and several parameters were measured, including body weight loss, SARS-CoV-2 RNA copies in the lungs, histopathological change in lung tissues, level of SARS-CoV-2 spikes in lung, levels of key inflammatory cytokines (IL-6, IL-1β, TNF-α) in lungs.

**Fig. 4.**
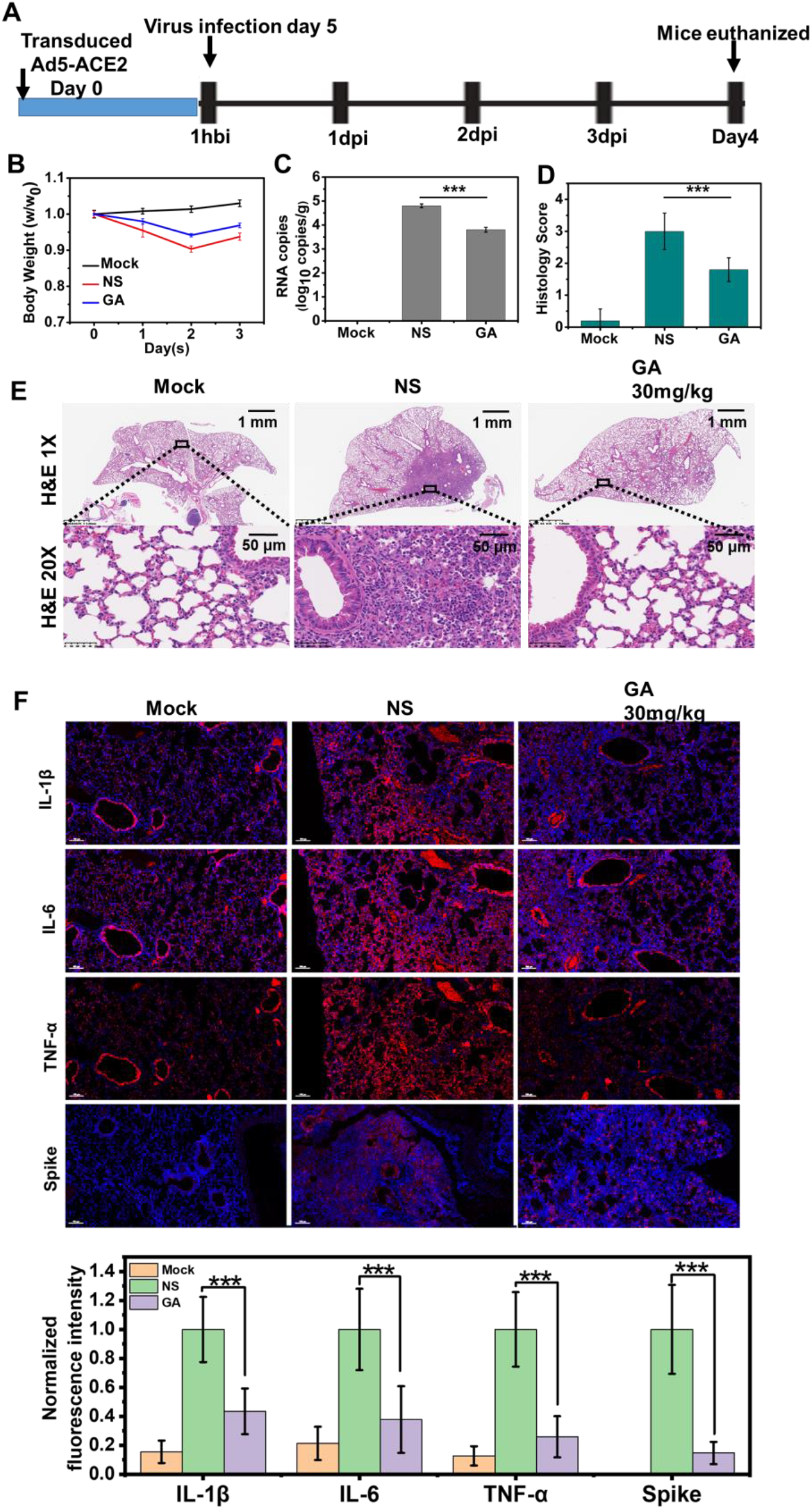
The GA inhibits virus replication, protects lung injury, and suppresses inflammatory cytokines expression in lungs of COVID-19 mice. (**A**) Schematic diagram of GA treated COVID-19 mice. (**B**) Weight loss of infected mice treated either with GA or NS. (**C**) Viral RNA copies in mouse lungs at day 4 post infection (unpaired t-test, ***p < 0.001). (**D**) Histopathological scores of lung injury in SARS-CoV-2 infected mice (unpaired t-test, ***p < 0.001). (**E**) Representative Hematoxylin-eosin (HE) staining of lungs from mice harvested at day 4 post infection, scale bar ~ 50 μm. **(F)** The fluorescence intensity of IL-6, IL-1β, TNF-α, and SARS-CoV-2 spike in lungs of NS and GA treated COVID-19 mice. Normalized fluorescence intensity of IL-6, IL-1β, TNF-α, and SARS-CoV-2 spike (unpaired t-test, ***p<0.001), scale bar ~ 50 μm.

As shown in **Figure 4**, infection of mouse model with SARS-CoV-2 resulted in a number of phenotypes, including obvious body weight loss, higher SARS-CoV-2 RNA copies in lung, and severe bronchopneumonia and interstitial pneumonia and infiltration of lymphocytes within alveolar. GA-treated COVID-19 mice are with better body weight in comparison of NS-treated COVID-19 mice (**Figure 4B**). We found that the average number of viral RNA copies in lung of GA-treated mice was ~10^4^, significantly lower than the average number found in NS-treated infected mice ~ 10^5^ (**Figure 4C**). The histopathological changes in mice lung tissues were assessed by grading the injuries in accordance with the International Harmonization of Nomenclature and Diagnostic Criteria (INHAND) scoring standard. As shown in **Figure 4D**, the GA-treated mice significantly reduced histopathological scores (~1.8) compared with that of NS-treated mice (~3.0). We further evaluated the therapeutic effects of GA using histopathological analysis of mouse lung tissues. As shown in **Figure 4E**, mice infected with SARS-CoV-2 showed severe lung inflammation following treatment with NS, the alveolar septum, bronchus, bronchioles and perivascular interstitium were significantly widened, along with an infiltration of higher numbers of lymphocytes and a small number of neutrophils. In addition, a small number of lymphocytes and exfoliated epithelial cells localized in the lumen of local bronchioles following NS treatment. Treatment with GA significantly abrogated lung inflammation in SARS-CoV-2 infected mice, the local alveolar septum, bronchi, bronchiole and perivascular interstitial widening significantly decreased. Although we still observed mild lymphocytic infiltration, the mucosal epithelium of bronchus and bronchioles was intact, and we failed to observe foreign bodies in the lumen in lung of GA treated mice.

Next, we measured SARS-CoV-2 spike and the key inflammatory cytokines in lung of the mice using immuno-fluorescent imaging. As shown in **Figure 4F**, the SARS-CoV-2 spike, IL-6, IL-1β, TNF-α expression level in the lung tissues of GA-treated infected mice were significantly lower than those found for NS-treated infected mice. Together, these results clearly demonstrated that GA inhibits virus replication, while also suppressing inflammatory cytokine expression, thus protecting the lungs of infected mice from inflammation injury.

### Via nasal dropping administration, GA shows better therapy outcome than Remdesivir in COVID-19 golden hamster model

Remdesivir was approved to treat COVID-19 in clinical now. In this study, GA is used to compare with Remdesivir to see which one is with better outcome in COVID-19 hamster treatment. A golden Syrian hamster model was generated according to recently published paper (21). Briefly, golden hamsters were randomly divided into five groups, and Hamsters were then infected intranasally using SARS-CoV-2 (1×10^5^ PFU) in a total volume of 50 μL DMEM (**SI 10**). One hour after SARS-CoV-2 infection, hamsters in NS group received normal saline (0.9% NaCl), hamsters in Remdesivir group received a dose of 25mg/kg.bw, and hamsters in first GA group received a dose of 10mg/kg.bw and in second GA group received a dose of 20mg/kg.bw. Mock group are mock-infected rats. All hamsters are treated by intranasally dropping administration. After virus infection at day 0, hamsters received GA or Remdesivir or NS treatment for further 3 times as shown in **Figure 5A**. All hamsters were euthanized at day 4 post infection, and several parameters were measured, including body weight loss, SARS-CoV-2 RNA copies in the lungs, pathological change in lung tissues, level of SARS-CoV-2 spikes in lung, and levels of key inflammatory cytokines (IL-6, IL-1β, TNF-α) in lungs.

**Fig. 5.**
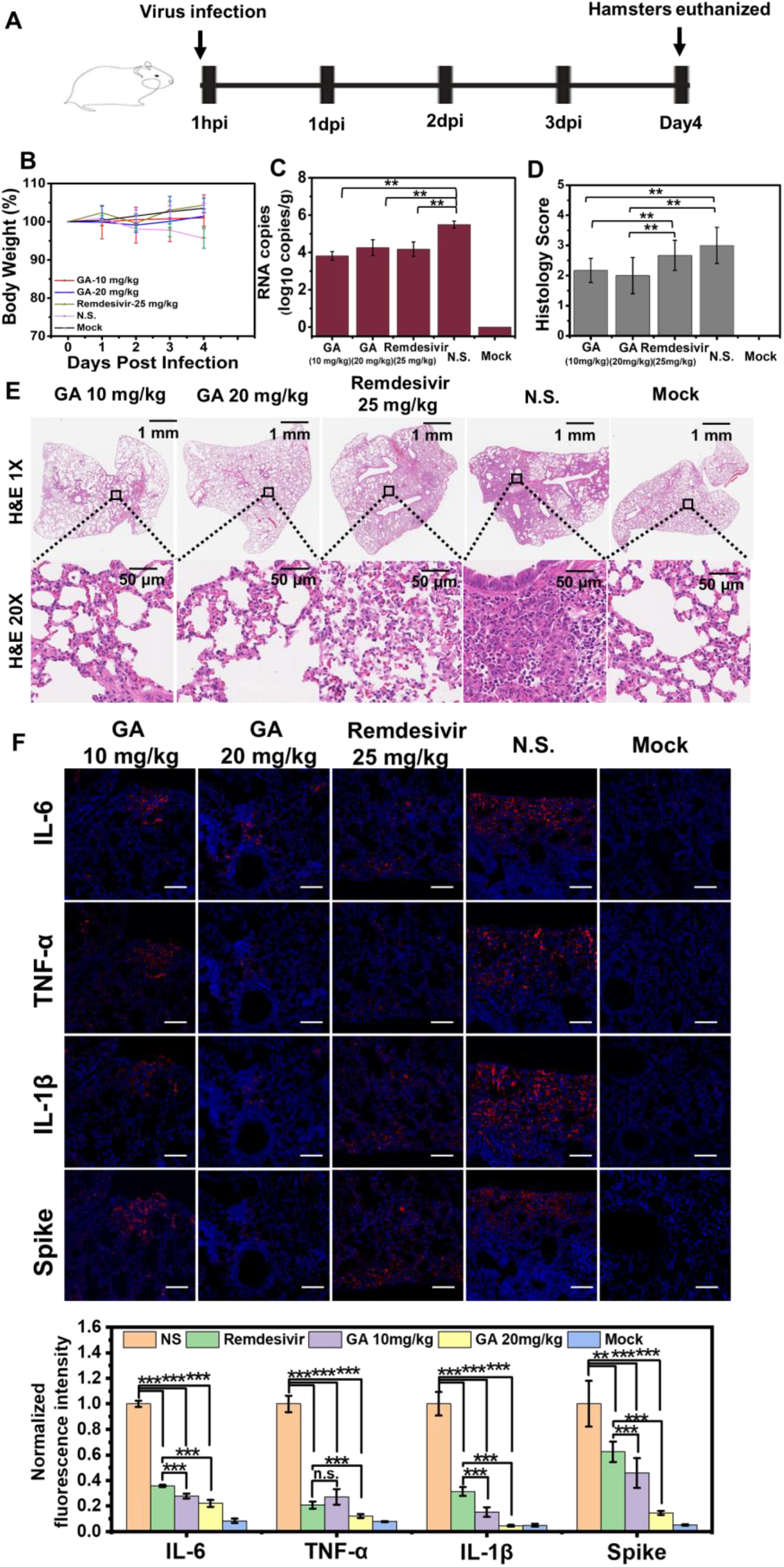
The GA and Remdesivir inhibit virus replication, protect lung injury, and suppresses inflammatory cytokines expression in lungs of COVID-19 golden hamsters. (**A**) Schematic diagram of GA or Remdesivir treated COVID-19 hamsters. (**B**) Weight loss of infected hamsters treated with GA or Remdesivir. (**C**) Viral RNA copies in hamster lungs at day 4 post infection (unpaired t-test, **p < 0.01). (**D**) Histopathological scores of lung injury of GA or Remdesivir treated SARS-CoV-2 infected hamsters (unpaired t-test, *p < 0.05, **p < 0.01). (**E**) Representative Hematoxylin-eosin (HE) staining of lungs from hamster harvested at day 4 post infection, scale bar ~ 50 μm. **(F)** The fluorescence intensity of IL-6, IL-1β, TNF-α, and SARS-CoV-2 spike distributed in lungs of GA or Remdesivir treated COVID-19 hamster. Normalized fluorescence intensity of IL-6, IL-1β, TNF-α, and SARS-CoV-2 spike (unpaired t-test, ***p<0.001, **p<0.01), scale bar ~ 50 μm..

GA or Remdesivir treated COVID-19 hamsters are with no significant body weight loss when compared to Mock hamsters, although NS-treated COVID-19 hamster obviously lost body weight (**Figure 5B**). We found that the average number of viral RNA copies in lung of GA (~10^3.8^ in 10mg/kg.bw group and ~10^4.2^ in 20mg/kg.bw group) or Remdesivir (~10^4.1^) treated hamsters was significantly lower than that found in NS-treated infected hamsters (~10^5.5^), GA and Remdesevir were in similar level in inhibiting virus replication in lungs of hamsters (**Figure 5C**).The histopathological changes in lung tissues are key index to assess the therapy effects of GA or Remdesivir in COVID-19 golden hamster. The lungs of hamsters were assessed by grading the injuries in accordance with the International Harmonization of Nomenclature and Diagnostic Criteria (INHAND) scoring standard. As shown in **Figure 5D and 5E**, the average histopathological score of virus-infected hamsters in NS group was approx. 3, the alveolar septum, bronchus, and perivascular interstitium were significantly widened, along with an infiltration of lymphocytes and neutrophils. For hamsters infected by SARS-CoV-2, treatment of Remdesivir in dose of 25mg/kg.bw got histopathological scores approx. 2.7, briefly the alveolar septum, bronchus, and perivascular interstitium were obviously widened, along with an infiltration of some of lymphocytes and neutrophils. Treatment of GA in dose of 10mg/kg.bw got pathological scores, approx. 2.3, and treatment with GA in dose of 20mg/kg.bw pathological score was approx. 2.0. GA treatment significantly decreased lung injury in comparison of Remdesivir treated SARS-CoV-2 infected hamsters. The local alveolar septum, bronchi, and perivascular interstitial widening were significantly decreased, along with an infiltration of smaller numbers of lymphocytes and neutrophils. According histopathological results, GA in dose of 10 mg/kg.bw or 20mg/kg.bw is with better therapy outcome than Remdesivir in dose of 25mg/kg.bw.

Next, we measured the level of SARS-CoV-2 spikes and inflammatory cytokines in lung of the hamsters using immuno-fluorescent imaging. As shown in **Figure 5F**, the SARS-CoV-2 spike expression level in GA and Remdesivir treated COVID-19 hamster was significantly lower than those found in NS group, and the spike level in lung of Remdesivir group is significantly higher than that of GA group. For inflammatory cytokine level in lung of virus infected hamsters, IL-6, IL-1β, TNF-α of Remdesivir or GA-treated rat were significantly lower than those found for NS-treated hamsters, and the IL-6, IL-1β, TNF-α in lung of Remdesivir group is significantly higher than that of GA group. As SARS-CoV-2 RNA copies in the lungs of GA or Remdesivir treated COVID-19 hamsters are in the same level, these immune-imaging results clearly demonstrated that GA intrinsically suppresses inflammatory cytokine expression in lungs of COVID-19 hamsters, the GA is with better outcome in suppression inflammatory cytokines expression when compared with Remdesivir.

### The tissue distribution of Au and histopathological studies of GA treated mice and golden hamsters

BALB/c mice (30mg GA/kg.bw via intraperitoneal injection) and golden hamster (20mg GA/kg.bw via intranasally dropping administration) were treated as description (**SI 11**). GA treatment resulted in none measurable side effects. Neither movement, out-looking, sleeping, nor eating behavior appeared to be affected. When we stained tissue sections with Hematoxylin-eosin (HE), we could not detect any pathological changes in GA-treated mice or GA-treated hamsters in **Figure S8**, indicating that our treatment dose of GA was safe. Next, we analyzed Au element distribution in lung, brain, liver, spleen, heart, and kidney using an ICP-MASS approach. As shown in **Table S2 and S3**. The distribution of gold in lungs, hearts, livers, kidneys, brains, and spleens can be beneficial for COVID-19 treatment, as it potentially inhibits SARS-CoV-2 replication and direct suppressing the expression of inflammatory cytokines therein (15, 16, 17). The Au element concentration in the lung of mice (intraperitoneal injection) and hamster (intranasally dropping) was approx. ~ 78.14 μg/g and ~ 416.12 μg/g, respectively. This result showed that GA is well adsorbed by lung after intranasally dropping method, which is a good way to deliver GA into lung for inhibit virus replication and directly suppress inflammation injury therein.

## Conclusion

In summary, our findings show that GA represents a promising therapeutic compound that combines intrinsically suppression of inflammatory cytokines expression with directly inhibition of SARS-CoV-2 replication in COVID-19 transgenic mice and golden hamsters. Via intranasally dropping, GA (10mg/kg.bw or 20mg/kg.bw) and Remdesivir (25mg/kg.bw) has similar inhibition ratio in lung viral load for COVID-19 hamsters. According histopathological studies, GA significantly suppress the lung injury when compared with Remdesivir in COVID-19 hamsters. According immuno-fluorescent observations, GA significantly suppress the lung inflammatory cytokines expression, this is attributed to GA directly inactivate NFkB pathway and further reduce inflammatory cytokines expression level in lung. The key factor that need to be considered when designing a drug for COVID-19 treatment is its toxicity. Our study showed that GA is characterized by very mild cytotoxicity and very low mice/rat acute toxicity. These features render GA an ideal candidate for safe and effective COVID-19 treatment.

## Acknowledgments

We thank Dr. Feng Wang (Wuxi Biortus Bioscience Co. Ltd, 6 Dongsheng West Road, Jiangyin 214437, China) provide the gift plasmid for expression SARS-CoV-2 M^pro^. We thank Dr. Yuanyuan Chen from Institute of Biophysics, Chinese Academy of Sciences for her technical support with the SPR assay. We thank the staffs from beamlines BL17U1, BL18U1 and BL19U1 at the Shanghai Synchrotron Radiation Facility for X-ray data collection. This work was supported by grants from National Natural Science Foundation of China (21727817, U2067214, 11621505). Author contributions: X. G. conceived the project; H. Z., F. Y., C. Z., Z. D., W. Z., Y. G., Y. Q., F. G. performed cell studies and enzyme activity; Y. G., J. F., B. H., P. C., B. S. performed the sample preparation, characterization and data collection; L. Z., X. G., X. G. performed theory calculation; Y. G., Y. D., J. Q., and H. J. processed data analysis; W.T., F. Y., J. Z. performed animal studies; X. G wrote the manuscript; all authors discussed and commented on the results and the manuscript.

## Data and materials availability

The atomic coordinates of the SARS-CoV-2 M^pro^ crystal structures in the AF incubating form, the GA incubating form, and the native form has been deposited in the Protein Data Bank under accession code 7DAT, 7DAU, and **7**DAV, respectively.

